# Effects of size and elasticity on the relation between flow velocity and wall shear stress in side-wall aneurysms: A lattice Boltzmann-based computer simulation study

**DOI:** 10.1101/711564

**Authors:** Haifeng Wang, Timm Krüger, Fathollah Varnik

**Author notes:** (HW); (FV).

## Abstract

Blood flow in an artery is a fluid-structure interaction problem. It is widely accepted that aneurysm formation, enlargement and failure are associated with wall shear stress (WSS) which is exerted by flowing blood on the aneurysmal wall. To date, the combined effect of aneurysm size and wall elasticity on intra-aneurysm (IA) flow characteristics, particularly in the case of side-wall aneurysms, is poorly understood. Here we propose a model of three-dimensional viscous flow in a compliant artery containing an aneurysm by employing the immersed boundary-lattice Boltzmann-finite element method. This model allows to adequately account for the elastic deformation of both the blood vessel and aneurysm walls. Using this model, we perform a detailed investigation of the flow through aneurysm under different conditions with a focus on the parameters which may influence the wall shear stress. Most importantly, it is shown in this work that the use of flow velocity as a proxy for wall shear stress is well justified only in those sections of the vessel which are close to the ideal cylindrical geometry. Within the aneurysm domain, however, the correlation between wall shear stress and flow velocity is largely lost due to the complexity of the geometry and the resulting flow pattern. Moreover, the correlations weaken further with the phase shift between flow velocity and transmural pressure. These findings have important implications for medical applications since wall shear stress is believed to play a crucial role in aneurysm rupture.

## Introduction

Brain aneurysms lead to almost 500,000 deaths per year [1]. Physiological flows, such as arterial blood flow, are generally characterized by the transport of fluids in compliant tubes [2], which involves complex fluid-structure interaction problems. The interplay of hemodynamics and elastic arterial vessel walls (e.g. through wall shear stress (WSS) and blood pressure) is believed to play a central role in the aneurysm initiation, growth and rupture [3–7].

It is known that abnormal WSS drives degradation of the vascular wall [8–10]. Flow velocity is one of the principal factors determining the magnitude of WSS. Compared with WSS, in practice, flow velocity is easier to be measured and often acts as a proxy observable for WSS [11–13]. In this work, we study how size, elastic deformation of the aneurysm and the waveform of transmural pressure determine the connection between flow velocity and wall shear stress as an important hemodynamic factor.

Aneurysms can be classified into two major categories: fusiform and saccular aneurysms; the latter is the most common type of aneurysm and the type most prone to rupture [14]. The saccular aneurysm has two subtypes: end-wall and side-wall aneurysms [15].

Intra-aneurysm (IA) hemodynamics is sensitive to morphological factors such as shape and size of the aneurysm [16–19]. It has been reported that an increase of aneurysm size leads to a decrease of the average IA flow velocity, 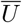, and wall shear stress, 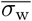, in the case of side-wall-type aneurysms [20, 21]. In these studies, the aneurysm wall is assumed to be rigid [20] or modeled as a stiff hyperelastic wall [21]. To date, however, the effect of wall elasticity and the resulting deformation is poorly studied.

Another objective of this work, therefore, is to investigate the combined effect of aneurysm size and wall softness on the variations of the average IA flow velocity and WSS, centering on the side-wall-type aneurysm.

We model the interaction of hemodynamics and wall elasticity in an idealized aneurysm using the lattice Boltzmann method (LBM) for the fluid flow, the finite element method (FEM) for the thin-walled vessel dynamics and the immersed boundary method (IBM) for the fluid-structure interaction.

The paper is organized as follows. The physical model is introduced in Sect Physical model. The numerical methods are presented in Sect Numerical methods, and benchmark tests are shown in Sect Validation. Simulation results using a curved artery with a side-wall aneurysm are presented and discussed in Sect Results and discussion, with a particular focus on the effect of aneurysm softness on IA hemodynamics. Our work is concluded in Sect Conclusion.

## Physical model

### Physical ingredients

Aneurysms occur in different sizes and shapes, and their detailed properties depend on the subject and progression state. According to [20, 21], the trends of the variations of flow velocity and wall shear stress due to the effect of aneurysm size are not specific to any particular side-wall aneurysm geometry. Therefore, after benchmark studies of flow through a straight cylindrical channel (Fig 1a), we use a representative model of a curved artery segment including a simple side-wall aneurysmal dome (Fig 1b and 1c). Without losing generality, this approach is similar to the simplification used in studies of the end-wall aneurysms [24–26].

**Fig 1.**
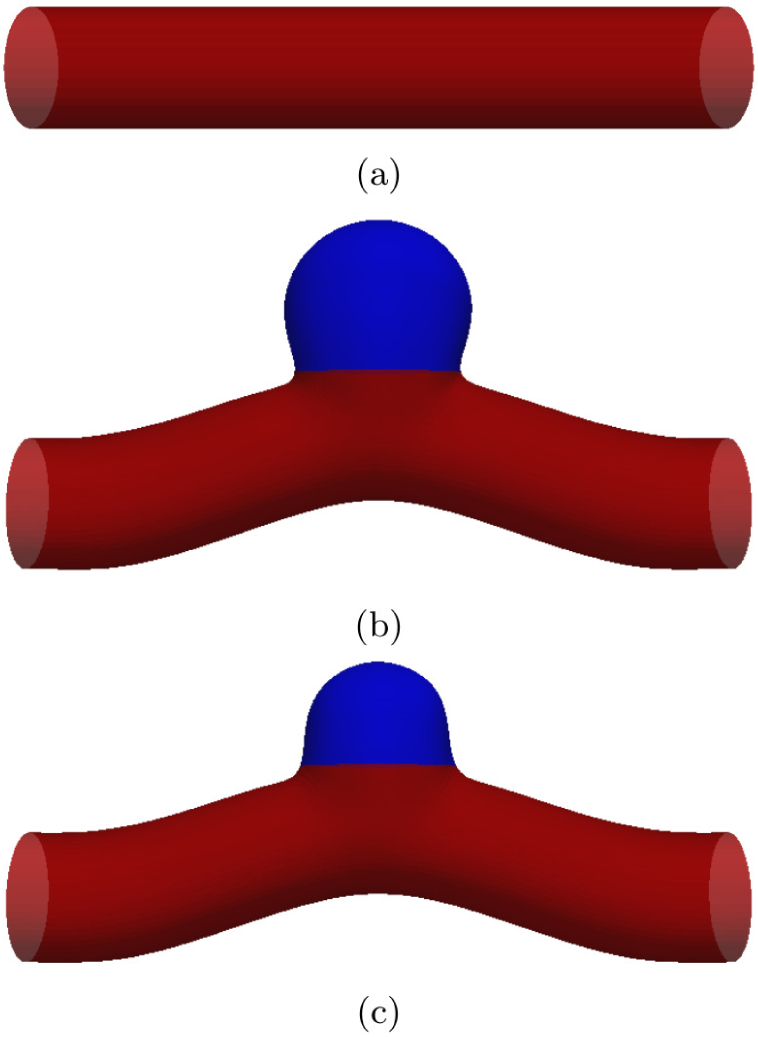
Geometries studied in this work. The straight circular tube in (a) is used to benchmark the code via comparisons with analytical solutions for both steady Poiseuille and pulsatile Womersley flows. Panels (b) and (c) show a curved blood vessel containing a side-wall aneurysm with different sizes, differing by a factor of 2.5 in aneurysm volume. Compared with the radius *R* of the parent vessel, the radii of the large (b) and small (c) aneurysm are approximately 1.5*R* and *R*, respectively. The domain colored in blue in (b) and (c) is modeled via a strain-softening Neo-Hookean law [22], while the remaining part of the vessel (red) obeys the strain-hardening Skalak law [23].

Vascular tissues are generally heterogeneous. Healthy arterial walls are normally strain-hardening due to the existence of collagen fiber constituents; aneurysms, however, often exhibit strain-softening behavior [27–30]. The arterial wall in our model is, therefore, divided into different regions (Fig 1b and 1c) with different constitutive laws. Similar to Charalambous et al. [31], we employ the strain-hardening Skalak (SK) model [23] for the healthy arterial wall:

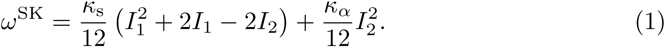

We use the strain-softening Neo-Hookean (NH) law (a special case of the Mooney-Rivlin law) [22] to model the aneurysm dome:

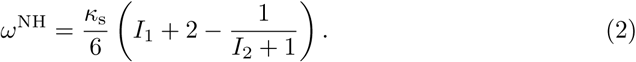

The SK and NH laws are described as areal energy densities *ω*^SK^ and *ω*^NH^, respectively. Both models have a surface elastic shear modulus *κ*_s_, and the SK model has an area dilation modulus *κ*_*α*_ (for the sake of simplicity, we choose *κ*_s_ = *κ*_*α*_ [31]). *I*_1_ and *I*_2_ are strain invariants depending on the local principal in-plane stretch ratios *λ*_1_ and *λ*_2_ as 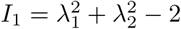 and 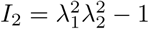.

The vessel wall is represented as a two-dimensional surface with zero thickness. However, one important effect closely related to a finite vessel wall thickness is resistance to bending. In order to account for this property in our membrane model, we explicitly introduce a local bending energy with a bending constant *κ*_b_, which can then give rise to forces normal to the wall (for further details, please refer to [32]).

The two free parameters *κ*_*α*_ and *κ*_b_ are related to the Young’s modulus *E*, wall thickness *d* and Poisson ratio *ν* according to [33]

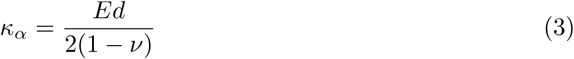

and

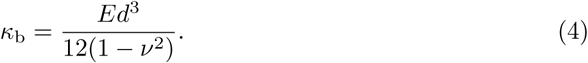

Thus, given the two material coefficients, *E* and *ν* and the thickness of the tissue, *d*, it is possible to determine the parameters of our model such that it reproduces the properties of a real vessel. This is explicitly shown below via a benchmark of the present study versus a 3D model. Obviously, the inverse mapping from a two-parameter model to a three-parameter one is not unique. This is not necessarily a drawback. Rather, it enhances the generic character of our model and allows the results obtained within these simulations to be transferred to vessels with different materials properties and wall thickness.

The transmural pressure *P*_tr_ is the pressure difference between the mean arterial pressure and the intracranial pressure. It is generally believed that, given heterogeneity of elastic properties along the arterial wall, *P*_tr_ is mainly responsible for the wall deformation and plays a crucial role in the progress of aneurysm growth as well as failure. Therefore, the transmural pressure needs to be taken into account in our model.

Although blood is non-Newtonian, it is often treated as an incompressible Newtonian fluid (e.g., [20, 21, 34–37]). Since the focus of this work is on the basic understanding of the effect of aneurysm size and softness rather than making accurate predictions, we model blood as a homogeneous Newtonian fluid with constant dynamic viscosity *η*.

Finally, we can assume the no-slip velocity boundary condition at the wall.

### Hemodynamic observables

Wall shear stress (WSS) is a crucial factor for the development of aneurysms. The fluid stress tensor includes pressure (isotropic stress) and viscous contributions (deviatoric stress):

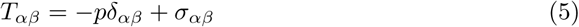

where *p* denotes pressure and ***σ*** the viscous stress tensor. The traction vector ***τ*** is the projection of the stress tensor onto the wall normal ***n***:

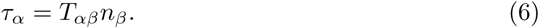

Here we are interested in the tangential (shear) component of ***τ*** whose magnitude is the WSS (*σ*_w_ = |***τ***^‖^|):

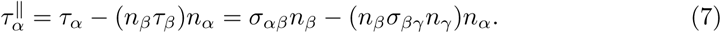

Note that the projection of the normal stress −*pδ*_*αβ*_ onto the tangential is zero, therefore *σ*_w_ is a function of ***σ***, but not a function of *p*.

For the aneurysm dome Ω (Fig 1b and 1c) with corresponding surface area *A* and volume *V*, we define the spatial WSS average

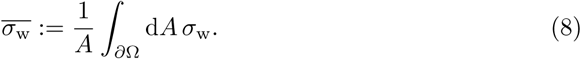

Similarly, the volume-averaged IA flow velocity is defined as

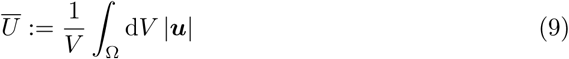

where ***u*** is the fluid velocity.

### Dimensionless groups

Based on the known material and flow properties (Young’s modulus *E*, artery radius *R*, transmural pressure *P*_tr_, arterial flow velocity *U* and fluid viscosity *η*), we can define a series of dimensionless groups characterizing the deformation of the arterial wall. The effect of transmural pressure can be described by

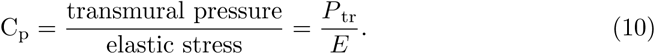

Viscous stresses may also contribute to the wall deformation as quantified by the capillary number

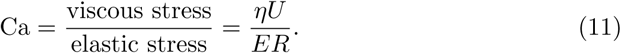

Finally, inertial deformation effects can be described by the Weber number

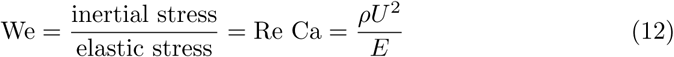

where Re = 2*ρUR/η* is the Reynolds number and *ρ* is the fluid density. Note that in normal arteries, *U* ≈ 0.3 m/s and *P*_tr_ ≈ 10 kPa, and, therefore, both viscous and inertial wall deformation effects are at least two orders of magnitude smaller than the transmural pressure contribution. This is in line with previous reports that the axial motion of vascular walls is negligibly small [38, 39]. We can therefore safely assume that deformation of the vessel wall is mainly due to the action of transmural pressure, rather than flow.

## Numerical methods

We employ a computational scheme combining the lattice Boltzmann method (LBM), finite element method (FEM) and immersed boundary method (IBM) [32] for a study of the coupled hemodynamic-artery problem outlined in Sect. Physical model. This hybrid method has previously been applied to suspensions of deformable red blood cells [40–42].

### Elastic wall model and force computation

The artery and aneurysm are modeled as thin elastic walls represented by a set of vertices and flat triangular facets. There are four force contributions acting on each vertex: the in-plane elastic shear force ***F***^s^, the normal bending force ***F***^b^, the transmural-pressure force ***F***^tp^ and a tether-like spring force ***F***^sp^.

The surface elastic shear energy *W*^s^ = ∫d*Aω*^s^ corresponding to the energy densities in Eq (1) or Eq (2) is computed via

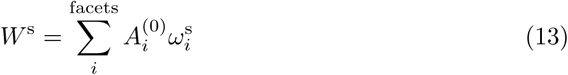

where *i* runs over all facets of the vessel wall, 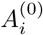 is the undeformed area of facet *i* and *ω*^s^ is either *ω*^SK^ or *ω*^NK^, depending on whether the facet *i* belongs to the healthy artery section or the aneurysmal dome.

The present model includes a bending resistance to avoid buckling of the arterial wall. The total bending energy of the artery is numerically approximated by

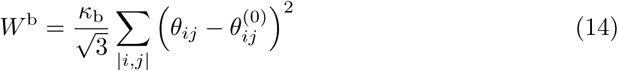

where *θ*_*ij*_ is the angle between neighbouring facet normals (i.e. facets *i* and *j* sharing two vertices), 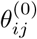 is the angle for the same pair of facets in the undeformed case and *κ*_b_ is the bending modulus [43].

Given the surface elastic shear energy *W*^s^ and the bending energy *W*^b^, the in-plane force ***F***^s^_*i*_ and bending force ***F***^b^_*i*_ acting on vertex *i* at position ***x***_*i*_ can be computed via the principle of virtual work:

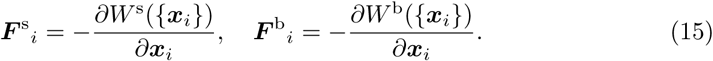

The transmural pressure *P*_tr_ is known to be one of the main factors dominating the enlargement and rupture of aneurysms [44–46]. There are basically two ways of modeling the transmural pressure. The most obvious one is to treat the transmural pressure as different pressures on either side of the arterial wall. This approach, however, poses a major challenge when combined with the immersed boundary method. We follow an alternative approach, introducing a transmural-pressure-based force ***F***^tp^ acting on each facet *i* along its normal direction:

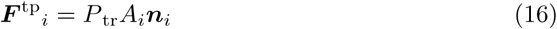

where *A*_*i*_ is the current area of facet *i* and ***n***_*i*_ the outward facing unit normal vector of the facet. ***F***^tp^ acts as a pressure drop across the wall without the necessity of having different pressures on either side of the wall in the numerical model. Note that ***F***^tp^ tends to dilate the artery if *P*_tr_ is positive. We emphasize that the transmural pressure has to be imposed, and it is generally a function of time: *P*_tr_(*t*). Fluid dynamics inside the deforming artery leads to additional spatio-temporal pressure fluctuations. This means that the *local* pressure drop across the arterial wall consists of a homogeneous term *P*_tr_(*t*) and a fluctuating term *P*_fluc_ which emerges from the simulation.

The fourth force contribution is a spring force ***F***^sp^ which is used to tether the ends of the artery:

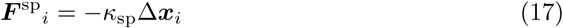

where *κ*_sp_ represents the spring constant, *i* denotes those vertices forming the open ends of the vessel segment and Δ***x***_*i*_ is the displacement of these vertices from their desired positions.

### Flow solver and fluid-structure interaction

In order to solve the incompressible Navier-Stokes equations, we employ a standard LBM on a D3Q19 lattice with the multiple-relaxation-time (MRT) collision operator for improved stability and accuracy [47–49]. Forces are included according to the Guo’s forcing scheme [50]. The viscous stress tensor is computed from the non-equilibrium populations according to [51].

Peskin’s IBM is used to couple the arterial wall and fluid dynamics [52]. Elastic forces calculated for each vertex *i* are spread to surrounding lattice nodes where they are used as input for the LBM. After the updated fluid velocity field has been obtained, fluid velocities are interpolated at the vertex positions via IBM. A tri-linear stencil with 2^3^ lattice nodes for each vertex is used during spreading and interpolation. The forward-Euler scheme is used to update all vertex positions. Further details can be found in [32].

### Interaction-related elastic energy

In addition to parameters such as maximum pressure difference along the blood vessel, which drives the flow, and size as well as flexibility of the aneurysm, the dynamics of the aneurysm wall may also strongly influence the IA flow field and thus lead to significant variations of the wall shear stress. In order to address this aspect, we will monitor the rate of variation of the elastic energy and investigate its correlations with WSS. The rate of elastic energy variation due to fluid-wall interactions can be calculated as

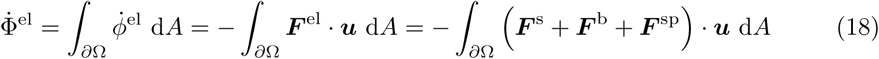

where the integration is performed over the entire aneurysm wall and 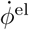 is the rate of change of the areal elastic energy density. For further reference and comparison, it is useful to also define a volume-averaged rate of elastic energy variation via 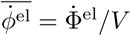.

## Validation

To validate the present model, we perform two different types of simulations: (i) steady and unsteady flow in a rigid tube and (ii) steady flow in an elastic tube.

### Steady and unsteady flows in rigid tube

We consider three-dimensional flow in a stiff tube with circular cross-section (Fig 1a). With the analytical solutions known for steady Poiseuille and pulsatile Womersley flows, this test allows a quantitative analysis of our model. All relevant simulation parameters are given in Table 1.

**Table 1.** Simulation parameters for Poiseuille and Womersley flow benchmarks. *P*_tr_: transmural pressure; *p*′: pressure gradient; *ρ*: fluid density; *η*: dynamic viscosity; *η*_B_: bulk viscosity; *R*: tube radius; *κ*_s_: elastic shear modulus; *κ*_*α*_: area dilation modulus; *κ*_b_: bending modulus; *κ*_sp_: spring modulus. Apart from the period *T*, parameters are the same in both flow tests. Using the maximum values of the velocities in the Poiseuille and Womersley flow, the Reynolds numbers are 178 and 125, respectively. The Womersley number is 2.48 in the case of pulsatile flow.

|  | Physical units | Lattice units |
| --- | --- | --- |
| $P_{\text{tr}}$ | 0 | 0 |
| $p'$ | 8.44 Pa/cm | $9.57 \cdot 10^{-8}$ |
| $T$ | 1 s | 224000 |
| $\rho$ | 1.055 g/cm<sup>3</sup> | 1.0 |
| $\eta$ | 4.22 mPa s | 0.0006 |
| $\eta_{\text{B}}$ | 11.7 mPa s | 0.1665 |
| $R$ | 2 mm | 11.459 |
| $\kappa_{\text{s}} = \kappa_{\alpha}$ | 13.6 N/m | 0.05 |
| $\kappa_{\text{b}}$ | $3.5 \cdot 10^{-8}$ N m | 0.004 |
| $\kappa_{\text{sp}}$ | 24 N/m | 0.09 |

A body force *F* along the axis of the tube is applied on all fluid nodes inside the tube to drive the flow. For Poiseuille flow, *F* is equivalent to the constant pressure gradient −*p*′ along the axis. For Womersley flow, the body force is given by *F* (*t*) = −*p*′ cos(*ωt*) where *ω* = 2*π/T* is the angular frequency and *T* is the sampling period.

The analytical Poiseuille solution is

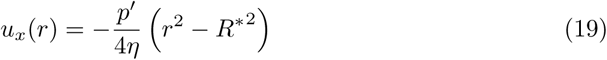

where *x* is the flow direction, *r* is the radial distance from the central axis, and *R*\* is the tube radius. For Womersley flow driven by the force *F* (*t*) above, the analytical solution is [53]

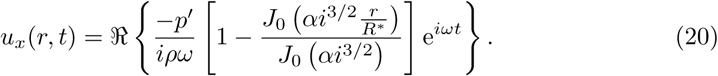

Here, ℜ{·} is the real part of a complex number, and *J*_0_(·) is the Bessel function of first kind and order zero. The Womersley number, which characterizes the oscillatory flow, is 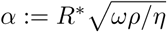. The viscous shear stress is 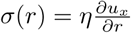. Note that 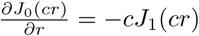, where *c* is a coefficient and *J*_1_(·) is the first-order Bessel function of the first kind. We use a Python script to solve the Womersley flow field.

The simulated flow velocities and shear stresses agree well with the corresponding analytical predictions in the two flow tests (Fig 2 and Fig 3). However, an extrapolation-based correction scheme is necessary to reach good agreement near the wall. The velocity and stress in direct wall vicinity are affected by the diffuse nature of the immersed boundary method. The basic idea behind the correction scheme is to enforce the no-slip condition in the diffuse interface IBM [54]. Specifically, fluid velocity and stress at the fluid nodes near the wall (i.e., at |*r/R*\*| = 11/11.035 in Fig 2 and Fig 3) are corrected using a second-order Lagrange extrapolation from corresponding information of their extrapolation points (i.e., at |*r/R*\*| = 10/11.035, 9/11.035 and 8/11.035). For general geometries, the extrapolation points may be off lattice lines. In such cases, fluid quantities (denoted by *f*) on each extrapolation node are computed by a linear interpolation of information at neighboring lattice nodes as *f* = (∑_*i*_ *f*_*i*_/*ℓ*_*i*_)/(∑_*i*_1/*ℓ*_*i*_) where *ℓ*_*i*_ indicates the distance between the extrapolation node and its neighboring lattice node *i*. For the sake of stability, when the extrapolation point is very close to a lattice node (e.g., *ℓ* < 0.0001), the value at that node is taken (*f* = *f*_*i*_). With this correction approach, the maximum relative L2 error of the shear stress *σ* is reduced from 31.2% to 2.2% in the Poiseuille flow and from 32.7% to 3.5% in the Womersley flow.

**Fig 2.**
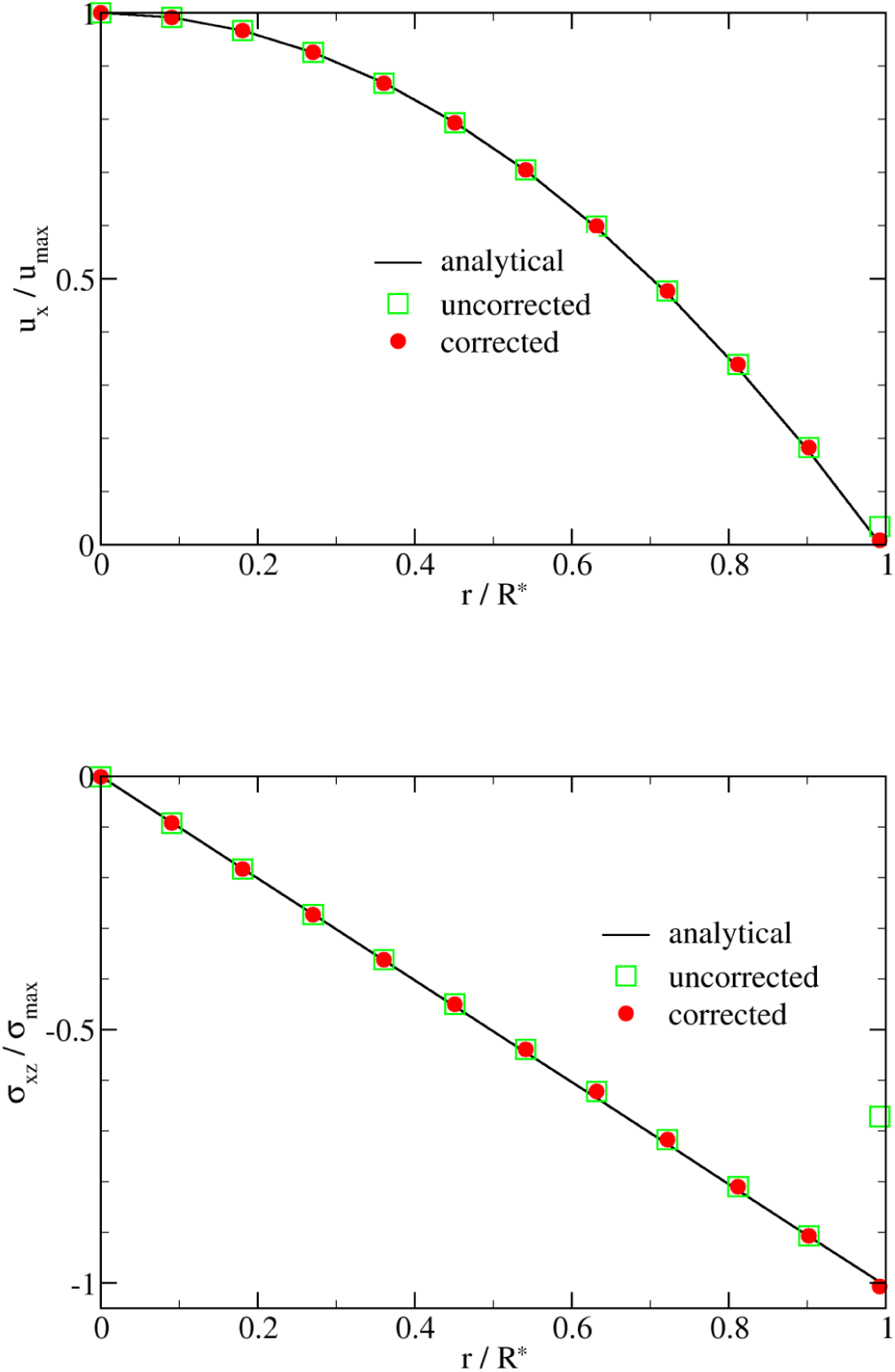
Poiseuille benchmark results. The simulation results (symbols) of fluid velocity *u*_*x*_ and shear stress *σ* are compared with the analytical steady solution of Poiseuille flow (black lines). Green squares and red circles, respectively, denote results without and with the extrapolation-based correction scheme. The data are taken along the *z*-coordinate axis. The difference is visible near the walls (*r/R*\* →1) and is caused by the diffuse nature of the immersed boundary method. In particular the shear stress benefits from the correction scheme.

**Fig 3.**
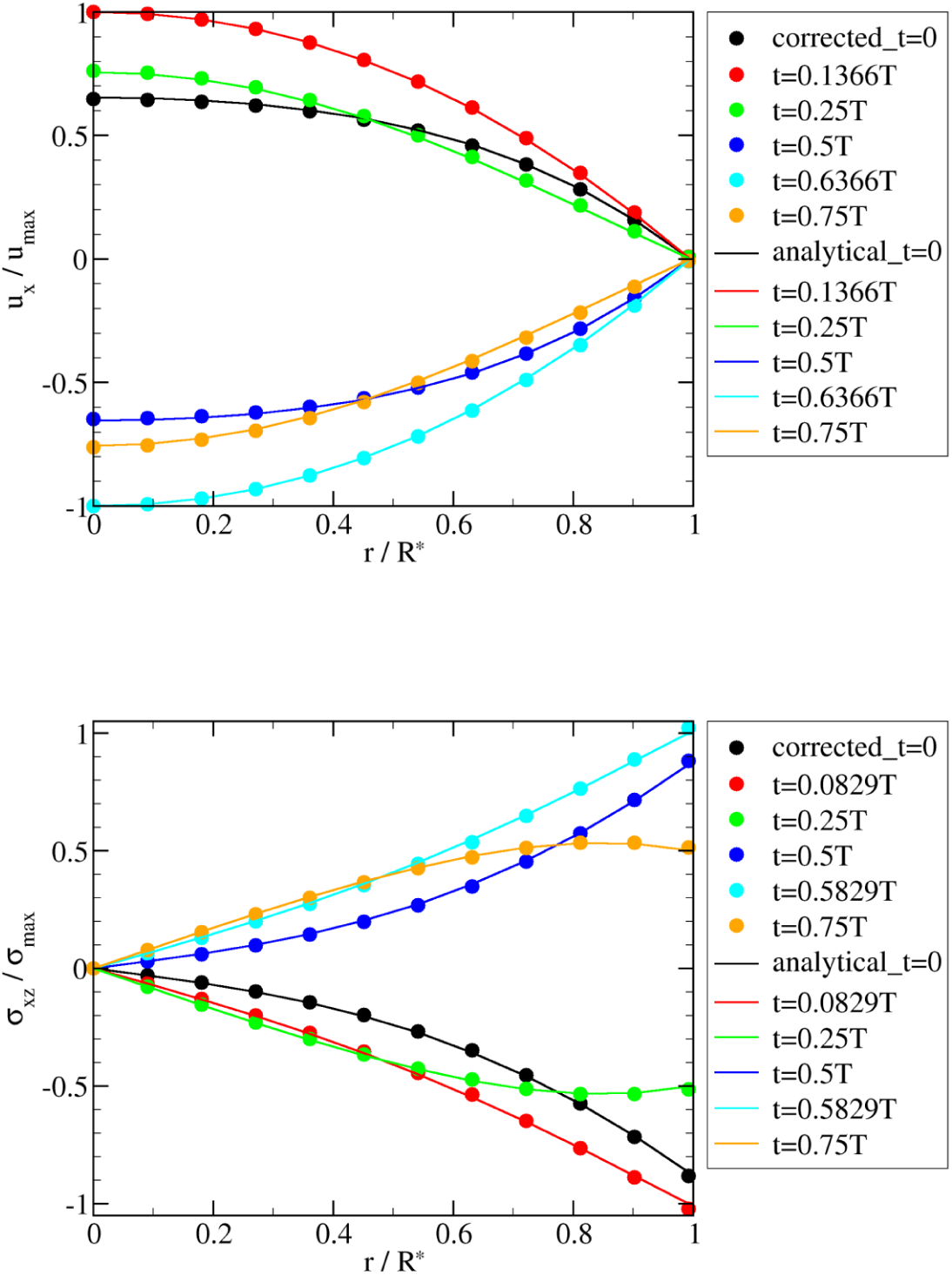
Womersley benchmark results. The simulation results (symbols) of fluid velocity *u*_*x*_ and shear stress *σ* are compared with the analytical solution of Womersley flow (lines). Only extrapolation-corrected results are shown. Each curve represents a different time step. Due to the correction scheme, the shear stress at the wall matches the analytical solution well.

Note that the wall location as recovered from the simulation is not exactly where it is expected to be. Due to the diffuse nature of the IBM, the location of zero velocity (as obtained from a quadratic extrapolation from the interior solution) is shifted with respect to the location of the Lagrangian markers. We treated the tube radius as fit parameter to obtain the best agreement with the simulated velocity profile. For our simulation parameters the apparent tube radius is *R*\* = 11.035Δ*x* compared to the input value of *R* = 11.459Δ*x*. Similar deviations have been reported in the literature [55–57] and are acceptably small.

### Steady flow in elastic tube

In order to show that the numerical model can faithfully reproduce flow in flexible three-dimensional geometries, we simulate steady flow in an elastic tube with a circular cross section (Fig 1a). In contrast to the steady Poiseuille flow, we consider the transmural pressure and the radial deformation of the tube walls. We use the well-developed software SimVascular [58] to generate reference results. SimVascular employs a simple linear elastic model for the walls.

To compare results from two different software tools, we need to ensure that the material models and properties used are identical. SimVascular requires the Young’s modulus *E*, the wall thickness *d*, and the Poisson ratio *ν* as input parameters. In the present model, the input parameters are the surface elastic shear modulus *κ*_s_, the area dilation modulus *κ*_*α*_, and the bending modulus *κ*_b_. Both sets of parameters are related according to Eq (3) and Eq (4). For the sake of stability of simulations with a certain resolution, small values of Young’s modulus *E* and transmural pressure *P*_tr_ are utilized. Nevertheless, it is C_p_, rather than *P*_tr_ or *E* alone, that dominates the wall deformation. Typical values used in SimVascular are *E* = 0.05 MPa, *d* = 0.15 mm and *ν* = 0.45. To match these parameters we set *κ*_s_ = 0.025, *κ*_*α*_ = 0.025 and *κ*_b_ = 0.002 in simulation units. The other simulation parameters are listed in Table 1. The length-diameter aspect ratio of the tube is 7.5. In all simulations, the ends of the tube are tethered and thus the deformation is not homogeneous along the tube axis; the radial displacements are zero at the ends and maximum in the middle region where the diametric strains (Δ*R/R*) are measured.

The combined effect of transmural pressure and elasticity on wall deformation is expressed by the dimensionless parameter C_p_, see Eq (10). We vary C_p_ by changing the transmural pressure *P*_tr_ in the present test case. Specifically, *P*_tr_ varies from 5.1 Pa to 153.2 Pa (i.e., 0.0001 ≤ C_p_ ≤ 0.003) in SimVascular simulations. Correspondingly, *P*_tr_ in our model ranges from 3.3 · 10^−6^ to 1 · 10^−4^ in simulation units. The radial strain Δ*R/R* of the wall is linearly proportional to C_p_ in SimVascular simulations due to the underlying linear elastic model. In order to investigate the strain-hardening behavior of the tube in the present model, five additional simulations with *P*_tr_ up to 6.5 · 10^−4^ in simulation units are carried out.

We find that simulation results using our model compare well with those obtained from SimVascular within the small strain regime (Fig 4a). In the large strain range, however, the strain-softening and strain-hardening features of the Neo-Hookean and Skalak models, respectively, become dominant and deviate from the behavior of the linear elastic model used in SimVascular.

**Fig 4.**
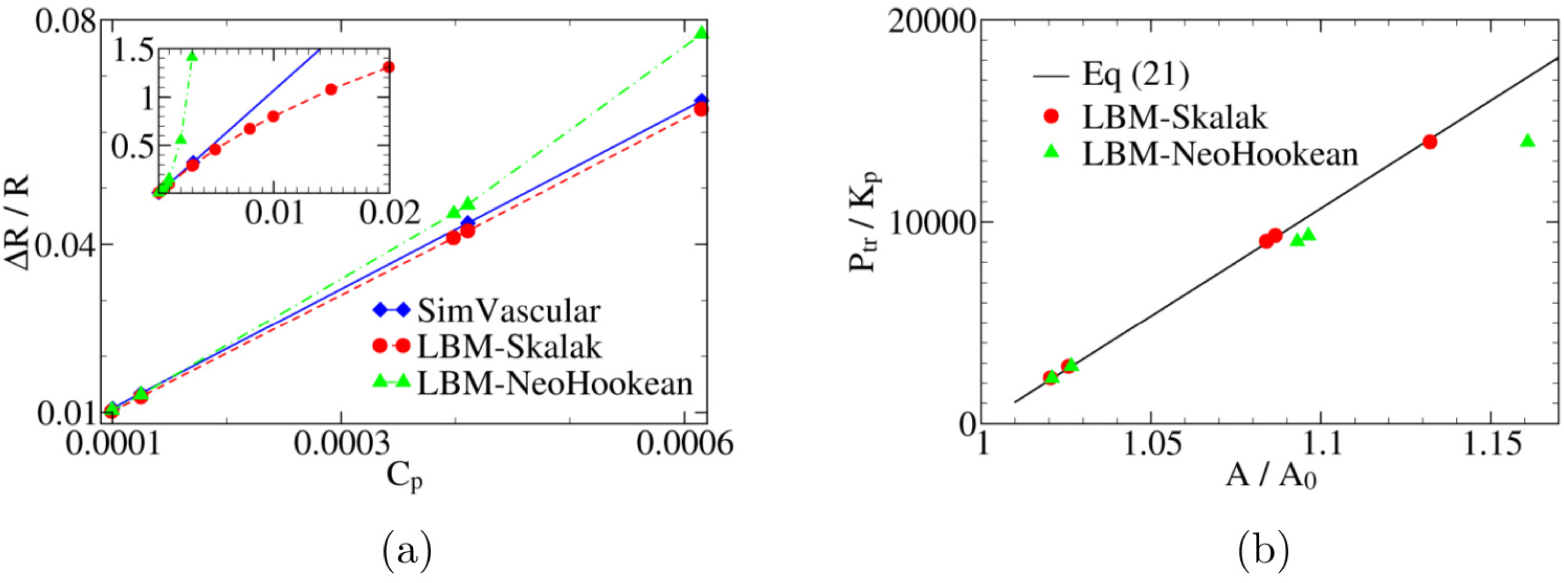
Benchmark of the elastic wall model. (a) The diametric strain, Δ*R/R*, versus C_p_ (the ratio of transmural pressure to Young’s modulus). SimVascular uses a linear elastic model and thus predicts Δ*R/R* ∝ C_p_. We use both the strain-hardening Skalak law and the strain-softening neo-Hookean model for the straight circular tube (Fig 1a). The parameters of these models are related via Eqs (3) and (4). As seen from the inset, in the limit of small strains (Δ*R/R* ≤ 0.1), these models produce the same linear behavior. (b) Validation of the model used in this work with the analytical result of reference [60]. The plot shows transmural pressure *P*_tr_ (normalized by *K*_*p*_) as a function of the cross-sectional area *A* (normalized by *A*_0_). As expected, simulation results using the Skalak model are in good agreement with the theoretical prediction, Eq (21).

Our simulation results also compare well with the so-called tube law (Fig 4b). In cases of wall expansion (*A/A*_0_ > 1), the tube law is [59, 60]

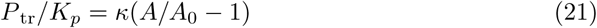

where, *K*_*p*_ is a constant proportional to bending resistance and can be expressed as *K*_*p*_ = *κ*_b_*/R*^3^ = *E*(*d/R*)^3^/[12(1 − *ν*^2^)] (see Eq (4)). *A*_0_ and *A* are the reference and deformed cross-sectional areas. The constant *κ* can be calculated as *κ* = 1.5(2*R/d*)^2^.

## Results and discussion

In the following, the combined effect of aneurysm size and wall softness on the flow characteristics within the *side-wall* aneurysm dome is investigated, to our best knowledge, for the first time. Brief introductions to both characteristic parameters and simulation setup are given before results are reported.

### Characteristic parameters

Blood and artery properties show variations among patients, but reference values based on a reasonable number of measurements have been reported. In general, the radius *R* of the cerebral artery ranges from 1.5 to 2.5 mm [21, 61, 62]. Young’s modulus *E* and Poisson’s ratio *ν* of the healthy cerebral artery are approximately 1–2 MPa and 0.45, respectively [21, 63]. The wall thickness is between 0.03 and 0.2 mm in most aneurysms, and is here assumed to be uniform and 0.15 mm, as adopted by [63]. There is no uniform ideal value for the transmural pressure *P*_tr_. However, its range lies roughly between 50 and 70 mmHg (i.e., between 6.7 and 9.3 kPa). Blood density *ρ* and dynamic viscosity *η* are within the range of 1–1.06 g/cm^3^ and 3–4 mPa s, respectively [21, 63]. The flow velocity *u* in cerebral arteries ranges from 0.1 to 1.0 m/s, and those in the common carotid and middle cerebral arteries are approximately 0.4 and 0.6 m/s, respectively [64–67].

In order to better highlight the effects arising from the interplay between elasticity, complex geometry and the time-dependent forces acting on the wall and on the fluid, it is useful to express physical quantities in such a way that trivial effects such as quadratic dependence of the maximum flow velocity and the linear dependence of maximum shear stress on tube radius as well as their linear dependence on the applied pressure gradient are “divided out”. A way to achieve this goal is to divide the quantities of interest such as flow velocity, *u*, wall shear stress, *σ*_w_, and viscous dissipation rate, 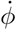, by their corresponding characteristic values in a straight cylindrical channel subject to a stationary Poiseuille flow, 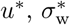 and 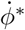. Throughout the paper, we will use the following reference values to make the corresponding quantities dimensionless: 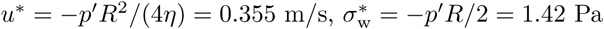, and 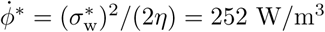. In this estimate, we used the values of the mean pressure gradient, *p*′ = 14.2 Pa/cm (Fig 5), vessel radius, *R* = 2 mm and dynamic blood viscosity, *η* = 4 · 10^−3^ Pa s.

**Fig 5.**
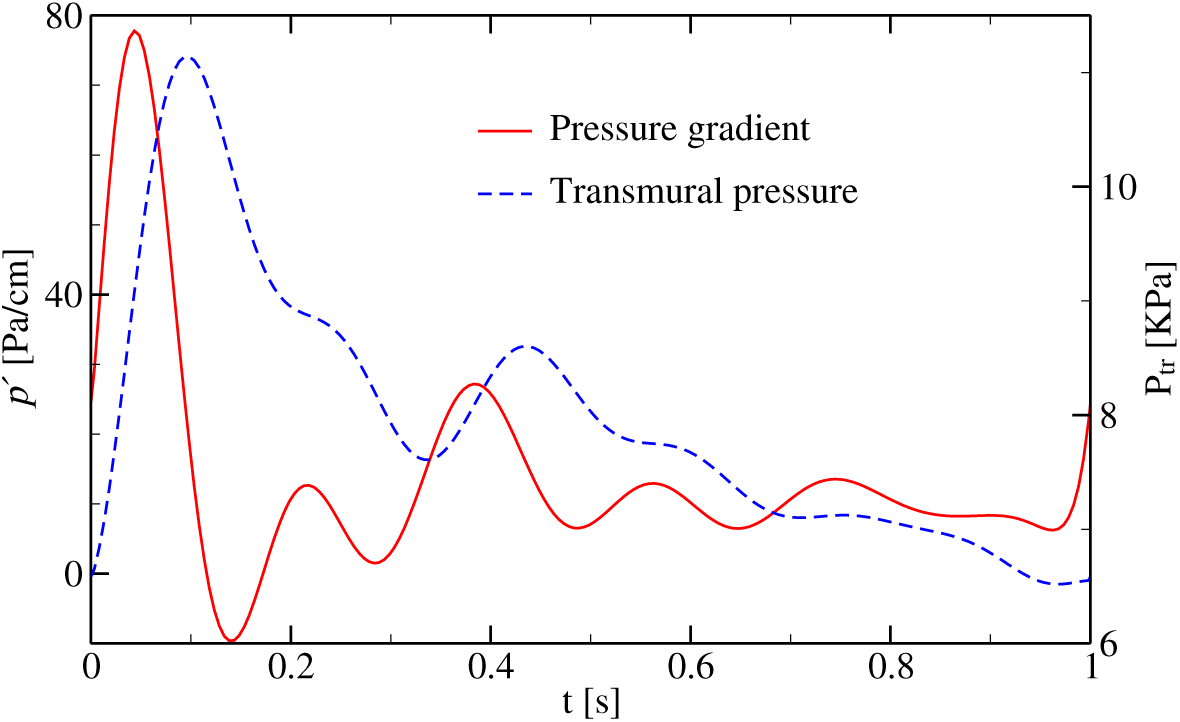
Imposed waveforms of pressure gradient, *p*′, and transmural pressure, *P*_tr_, over one period. The driving force is applied at the inlet as a force density, *f* = −*p*′. The transmural pressure is realized as a constant force per unit area of the wall acting in the direction of its normal vector. The profiles are taken from [46, 68].

### Simulation setup

We consider pulsatile flow in seven different cases:

1. Reference channel: straight cylindrical artery (Fig 1a) with elastic Young’s modulus *E*_0_ = 2 MPa. In the following, we will refer to this choice of *E*_0_ as the “Normal Elastic” (NE) case.
2. LA/stiff: large aneurysm (Fig 1b) with *P*_tr_ = 0
3. LA/NE: large aneurysm (Fig 1b) with normal elasticity *E*_0_
4. LA/RE: large aneurysm (Fig 1b) with reduced elasticity *E*_0_/3
5. SA/stiff: small aneurysm (Fig 1c) with *P*_tr_ = 0
6. SA/NE: small aneurysm (Fig 1c) with normal elasticity *E*_0_
7. SA/RE: small aneurysm (Fig 1c) with reduced elasticity *E*_0_/3

The initial (equilibrium) volume of the large aneurysm is approximately twice as large as that of the small aneurysm. The cases with zero transmural pressure mimic nearly rigid walls and allow to focus on the effects of size and deformability separately. All relevant parameter values used in the following simulations are given in Table 2.

**Table 2.** Simulation parameters for pulsatile flow in artery with/without aneurysm. *N*_*x*_, *N*_*y*_, *N*_*z*_: size of simulation box; see Fig 5 for waveforms of pressure gradient *p*′ and transmural pressure *P*_tr_; see Table 1 for other parameters. The values of *κ*_s_, *κ*_*α*_ and *κ*_b_ correspond to *E*_0_ = 2 MPa. For the softer aneurysm (*E*_0_/3), their values become three times smaller.

|  | Physical units | Lattice units |
| --- | --- | --- |
| $(N_x, N_y, N_z)$ | – | (128, 64, 100) |
| $T$ | 1 s | 427657 |
| $\rho$ | 1.055 g/cm<sup>3</sup> | 1.0 |
| $\eta$ | 4 mPa s | 0.0003 |
| $\eta_B$ | 22.2 mPa s | 0.1666 |
| $R$ | 2 mm | 11.634 |
| $\kappa_s = \kappa_\alpha$ | 272.7 N/m | 0.278 |
| $\kappa_b$ | $7.05 \cdot 10^{-7}$ N m | 0.024 |
| $\kappa_{sp}$ | 98 N/m | 0.1 |

Pulsatile flow is generated by applying a time-dependent body force (equivalent to the pressure gradient *p*′) at the inlet. The transmural pressure is applied on the entire wall. In reality, it is challenging to obtain both pressure gradient (or flow velocity) and transmural pressure waveforms from the same underlying mechanism (heart beat). Therefore, they are fed separately into the present model.

In practice, fluid velocity is often used as a proxy observable for wall shear stress [11–13]. While obvious in the case of a straight cylindrical channel, the complex shape of the aneurysm and its deformability introduce new aspects whose effects on the connection between flow velocity and wall shear stress needs a thorough investigation. Here, we address this issue via two different classes of spatially and temporally resolved simulations using the present hybrid model.

1. First we employ realistic waveforms (Fig 5) for both the pressure gradient and the transmural pressure. Given the chosen input parameters, the Reynolds, Womersley and C_p_ values inside the parent artery region with normal elasticity are approximately 350, 3 and 0.005, respectively. Within the aneurysm dome, the Reynolds number is almost 10 times smaller and C_p_ changes according to the variation of aneurysmal wall elasticity.
2. In the second class of simulations, to simplify the analysis, we assume sinusoidal waveforms for the pressure gradient, *p*′(*t*) = *p*′(0) sin(2*πt/T*), and the transmural pressure, *P*_tr_(*t*) = *P*_tr_(0) sin(*θ* + 2*πt/T*). We introduce a phase shift *θ* between both waveforms in order to quantify the individual contributions of both waveforms to the flow field.

Simulations are repeated for six cycles, which is sufficient to reach a time-periodic flow field. In all the simulations whose results are reported below, the open ends of the blood vessel are tethered. A body force (equivalent to pressure gradient *p*′ in Fig 5) is imposed at the inlet, and periodic boundary conditions are applied along the vessel axis.

### Realistic waveforms: effect of aneurysm softness

Figure 6 shows the time evolution of the space-averaged WSS and flow velocity in the straight artery (reference case). A comparison with the imposed waveforms in Fig 5 reveals that the time evolution of the hydrodynamic observables in the reference case basically follow the pressure gradient and transmural pressure. This is to be expected since the straight artery does not have complex geometrical features that could affect the flow. The average flow velocity and WSS are similar to those obtained for the corresponding steady Poiseuille flow.

**Fig 6.**
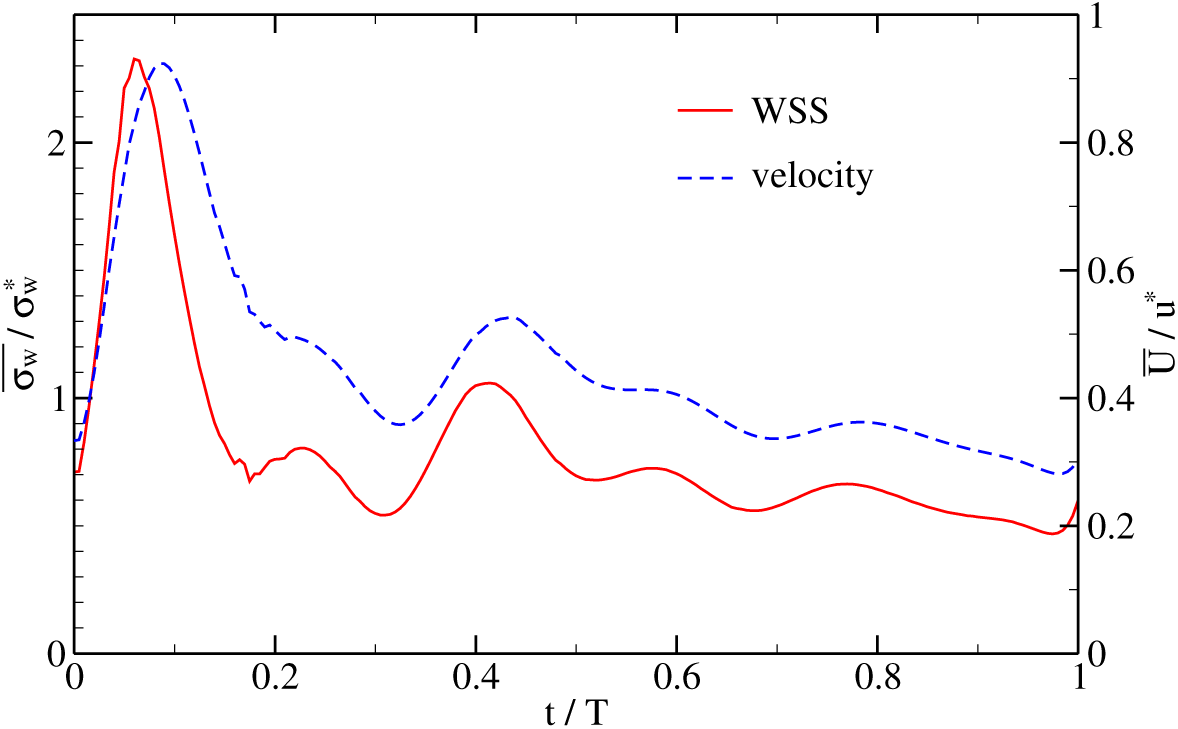
Space-averaged wall shear stress and flow velocity in a straight artery (reference case). Both WSS, 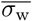, and flow velocity, 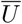, are averaged over a circular cross section in the midway between inlet and outlet and normalized by their reference values 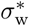 and *u*\*, respectively (page 11).

The time evolution of the intra-aneurysm (IA) WSS and flow velocity for the large (LA) and small (SA) aneurysm cases are shown in Fig 7. Similar to the straight tube case, the key features of the temporal variations of WSS and flow velocity stem from the pulsatility of flow rather than the existence of the aneurysm (compare Fig 5 and Fig 7). However, both the aneurysm size and wall softness impact the finer details of the IA hemodynamic characteristics as will be discussed in the following.

**Fig 7.**
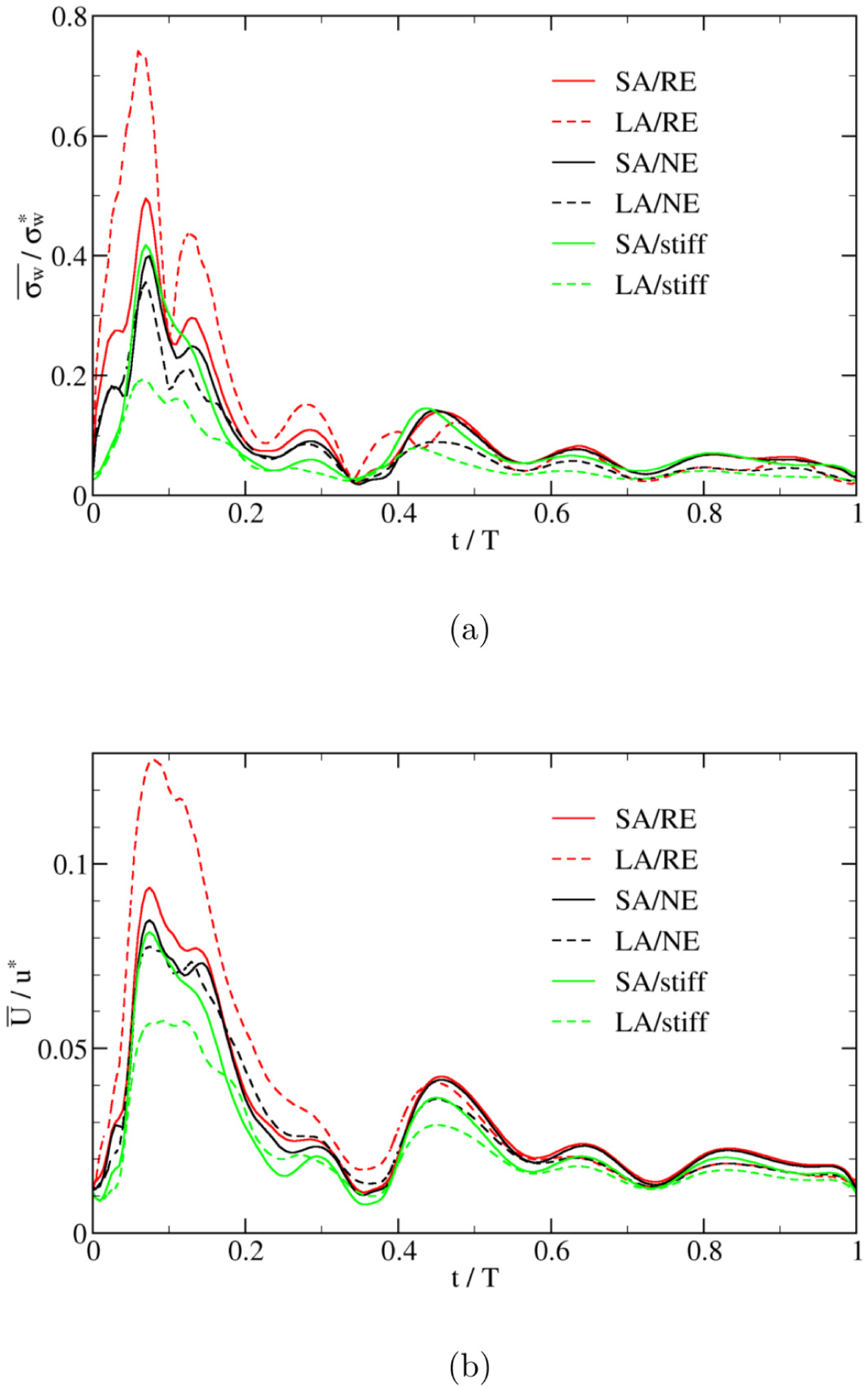
Time evolution of space-averaged wall shear stress and flow velocity inside the aneurysmal dome. Both (a) WSS, 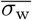, and (b) flow velocity, 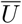, are normalized by their reference values in a straight cylindrical tube (page 11). Acronyms in the legends: SA=Small Aneurysm; LA=Large Aneurysm; NE=Normal Elasticity (using *E*_0_); RE=Reduced Elasticity (using *E*_0_/3) (page 12).

For the small-deformation cases (normal elasticity, LA/NE and SA/NE, and non-deformable, LA/stiff and SA/stiff), we find that the spatial average wall shear stress, 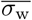, and flow velocity, 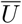, decrease with increasing the aneurysm size. In these cases the IA flow can be approximated by flow in a rigid cavity. The larger aneurysm has more potential to result in flow regions of recirculation and stagnation which are generally accompanied by lower WSS. This is in line with reported results in [20] and [21].

In real aneurysms, arterial wall stiffness may decrease due to tissue degradation. For the softer aneurysm (cases SA/RE and LA/RE), we find that 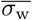 and 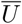 *increase* with aneurysm size (solid and dashed red lines in Fig 7). Note that the size of the initially larger aneurysm (LA) always remains larger than that of the smaller aneurysm. The variation of aneurysm volume (i.e., volumetric strain) directly follows the time evolution of the transmural pressure in Fig 5. The maximum volumetric strains are 61% and 86% in the cases SA/RE and LA/RE, respectively. Apparently, the aneurysm size no longer dominates the IA WSS and flow velocity behavior if the aneurysm is sufficiently soft. We can, therefore, conclude that there are at least two independent mechanisms controlling the IA flow properties: (i) size and (ii) deformability of the aneurysm dome.

Focusing on either the large or the small aneurysm, Fig 7 demonstrates that, the softer the aneurysm, the larger the IA WSS and the flow velocity, all averaged within the aneurysm domain. In order to gain more insight into the problem, a detailed view of the velocity field is shown in Fig 8 for three choices of a non-deformable (rigid) wall (indicated as stiff in the plot), a membrane with an intermediate (‘normal’) elastic constant, *E*_0_, and a soft aneurysm tissue with reduced elasticity. The plot shows that the flow velocity within the soft aneurysm reaches higher values than the two other cases. This observation may be rationalized as follows. Since the velocity of fluid at the wall is equal to that of the wall (stick boundary condition), a rigid and immobile vessel acts like a sink with respect to fluid momentum and thus decelerates the flow. A deformable tissue, on the other hand, can move under the action of transmural pressure and thus represents a less severe obstacle to the flow. In fact, depending on the phase of the flow with respect to the motion of the membrane, a partial enhancement of the fluid velocity is also possible. One possibility here would be that an elastic wall can store energy during one period of a cycle and release it during a different period again. This effect becomes stronger as the membrane softens further and its deformability increases.

**Fig 8.**
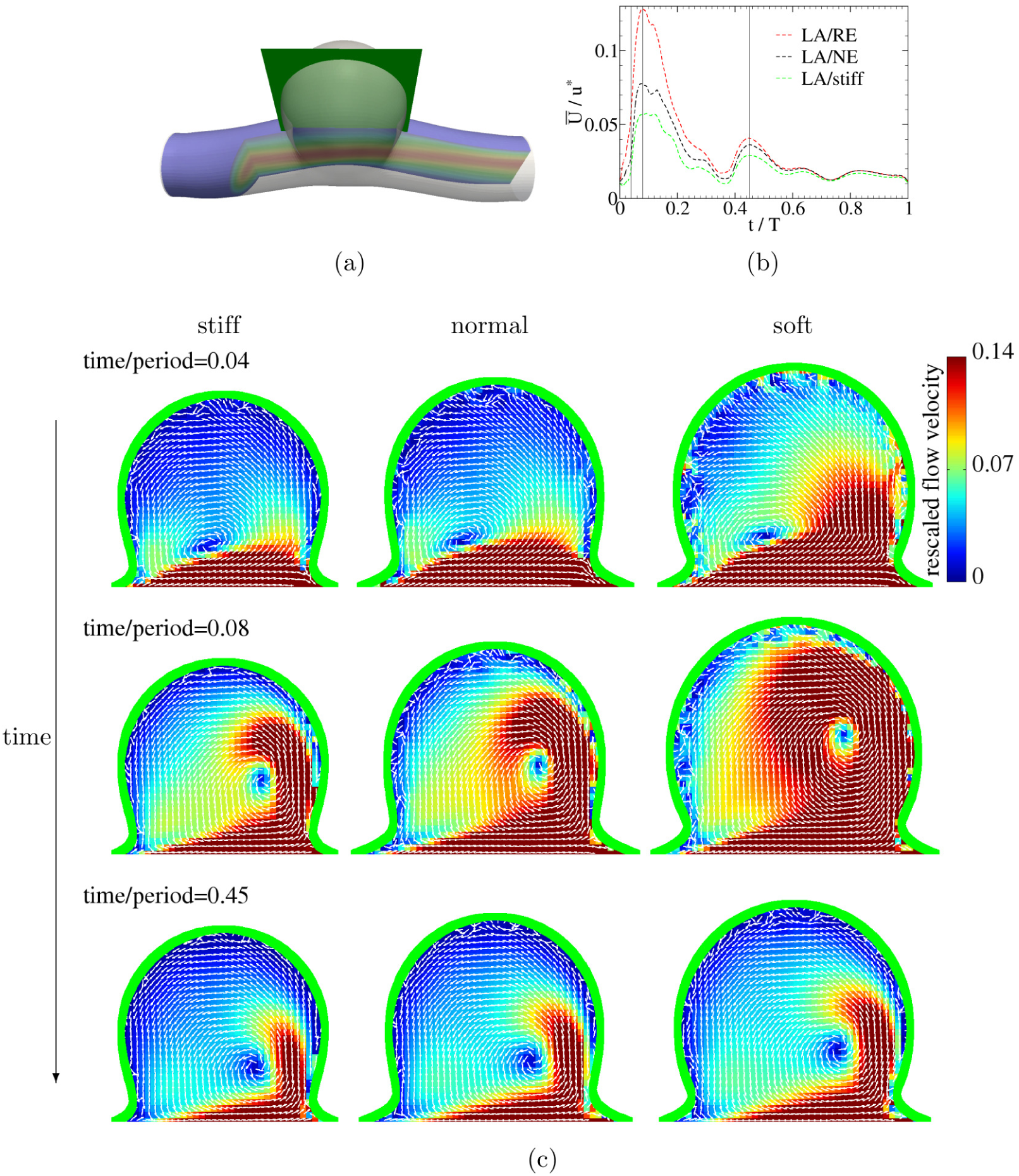
Flow field analysis. Upper row: (a) A view of the flow domain. The green plane shows the 2D cut through the aneurysm (Fig 1b) within which the flow field is visualized. (b) Average velocity (rescaled by *u*\* = 0.355 m/s) within the aneurysm for three different choices of wall elasticity. Acronyms refer to: LA=Large Aneurysm; NE=Normal Elasticity (using *E*_0_ as given on page 12); RE=Reduced Elasticity (using *E*_0_/3); stiff=non-deformable. The three lower rows illustrate the velocity field (again rescaled by *u*\* = 0.355 m/s) within aneurysm at different stages of a cycle as indicated by vertical lines in (b). White arrows indicate velocity vectors. Green line represents the aneurysm wall. It is clearly visible that the fastest flow develops within the softest aneurysm. An animation of the velocity field is available in S1 Appendix.

Such correlation between wall softness and flow parameters may, however, be reduced or even violated to some extent when effects of input waveforms become significant, as will be discussed in the following section.

From the viewpoint of energy, the effect of wall softness can be indicated by the variation of elastic energy. The elastic energy follows the transmural pressure (Fig 9a) since the shear and inertial contributions are relatively negligible (at least two orders of magnitude smaller) compared with the transmural-pressure contribution. As expected, the elastic contribution disappears in the case of a stiff wall (zero transmural pressure) (Fig 9a).

**Fig 9.**
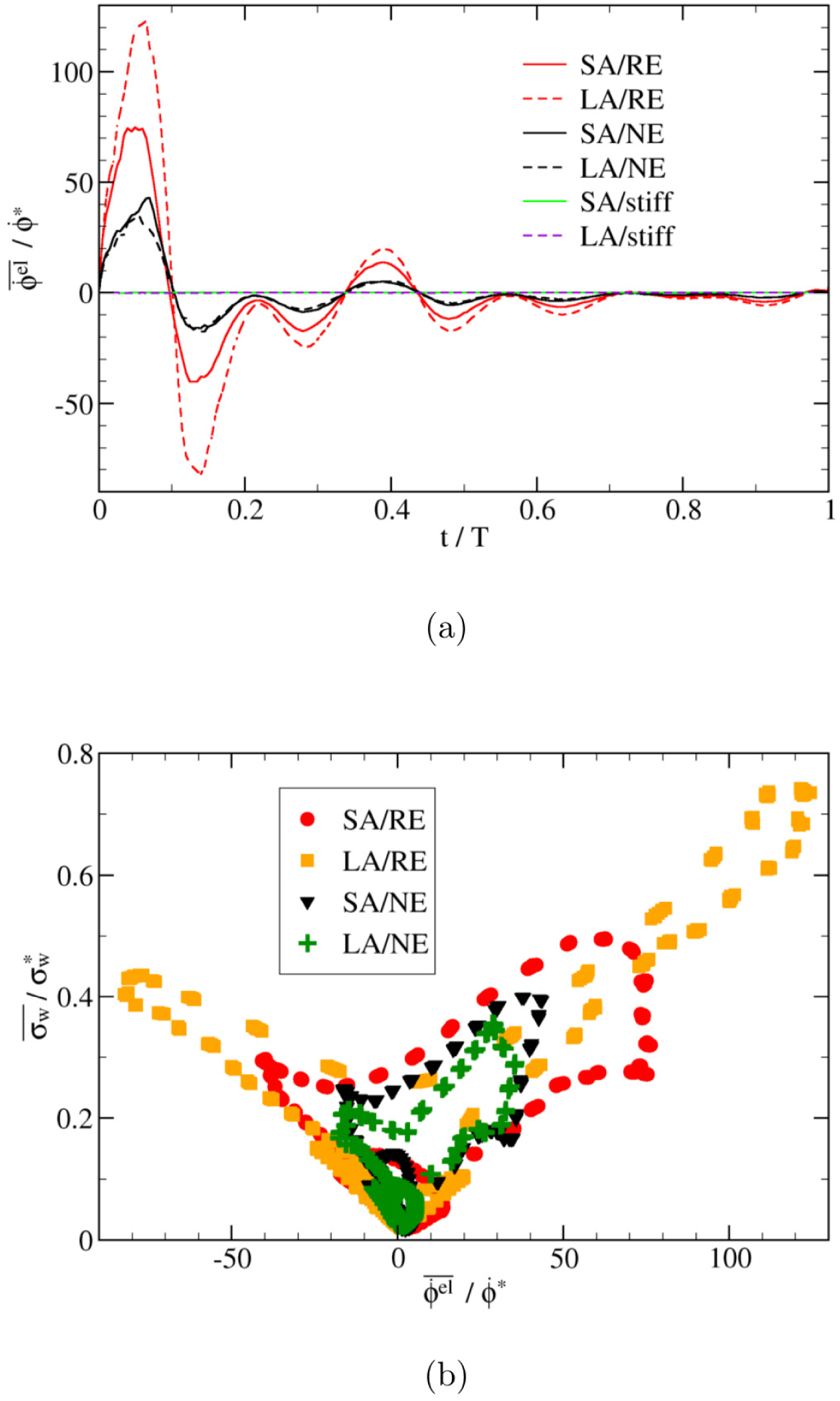
Elastic energy variation and correlation with wall shear stress. (a) Time evolution of volume-averaged variation rate of elastic energy, 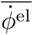, and (b) its relation to wall shear stress, 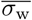, inside the aneurysmal dome. Both 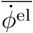 and 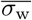 are computed as averages over the entire aneurysm region and are normalized by their reference values in a straight cylindrical tube (page 11). Acronyms in the legends: SA=Small Aneurysm; LA=Large Aneurysm; NE=Normal Elasticity (using *E*_0_); RE=Reduced Elasticity (using *E*_0_/3) (page 12).

In the case of side-wall aneurysms with realistic waveforms, we find a linear correlation between the rate of elastic energy variation and the wall shear stress (Fig 9b). No clear relation between the IA flow velocity and the rate of change of elastic energy has been found, though. Note, however, that it might be impossible to find a universal correlation between elastic energy rate and wall shear stress for all types of aneurysm geometries and input signals because the hemodynamics in aneurysms is sensitive to a number of conditions, including shape parameters such as inflow angle.

Naively, the IA WSS should follow the imposed pressure gradient. The deformation of the aneurysm, largely caused by the transmural pressure waveform, however, modifies the flow field in the aneurysm, which in turn affects the WSS. In order to investigate this effect, we consider sinusoidal input waveforms for the large aneurysm (Fig 1b) in the following.

### Sinusoidal waveforms: correlations between WSS and fluid velocity

Using sinusoidal waveforms for both pressure gradient *p*′ and transmural pressure *P*_tr_, together with a phase shift *θ* for the latter, we consider the following cases:

1. *θ* = −*π/*2: cosine-type *P*_tr_, i.e., *p*′ and *P*_tr_ are uncorrelated
2. *θ* = 0: sine-type *P*_tr_, i.e., *p*′ and *P*_tr_ are positively correlated
3. *θ* = *π*: sine-type *P*_tr_ with inverse amplitude, i.e., *p*′ and *P*_tr_ are negatively correlated

We begin with an analysis of the flow in the parent vessel before moving on to the aneurysmal dome. Well away from the aneurysm, flow velocity and WSS are governed by the input pressure gradient, and the phase shift between this driving force and the transmural pressure does not play any significant role (Fig 10). The cross-sectional average values of the flow velocity and wall shear stress are linearly correlated in the parent vessel (Fig 11a). There also exists a linear correlation between maximum velocity and average WSS and between maximum velocity and maximum WSS (the corresponding data is very similar to Fig 11a and is thus not shown). This strong correlation is also reflected in a high Pearson correlation coefficient, which turns out to be close to unity outside the aneurysm domain for all the investigated cases: 0.98, 0.99, 0.99 for *θ* = 0, −*π/*2 and *π*, respectively. Other combinations of size and transmural pressure give similarly large correlation coefficients: 0.99 (LA/NE/*θ* = *π*), 0.96 (LA/stiff) and 0.98 (SA/stiff). For the fluid outside the aneurysm, therefore, one can safely use the flow velocity as a proxy for wall shear stress.

**Fig 10.**
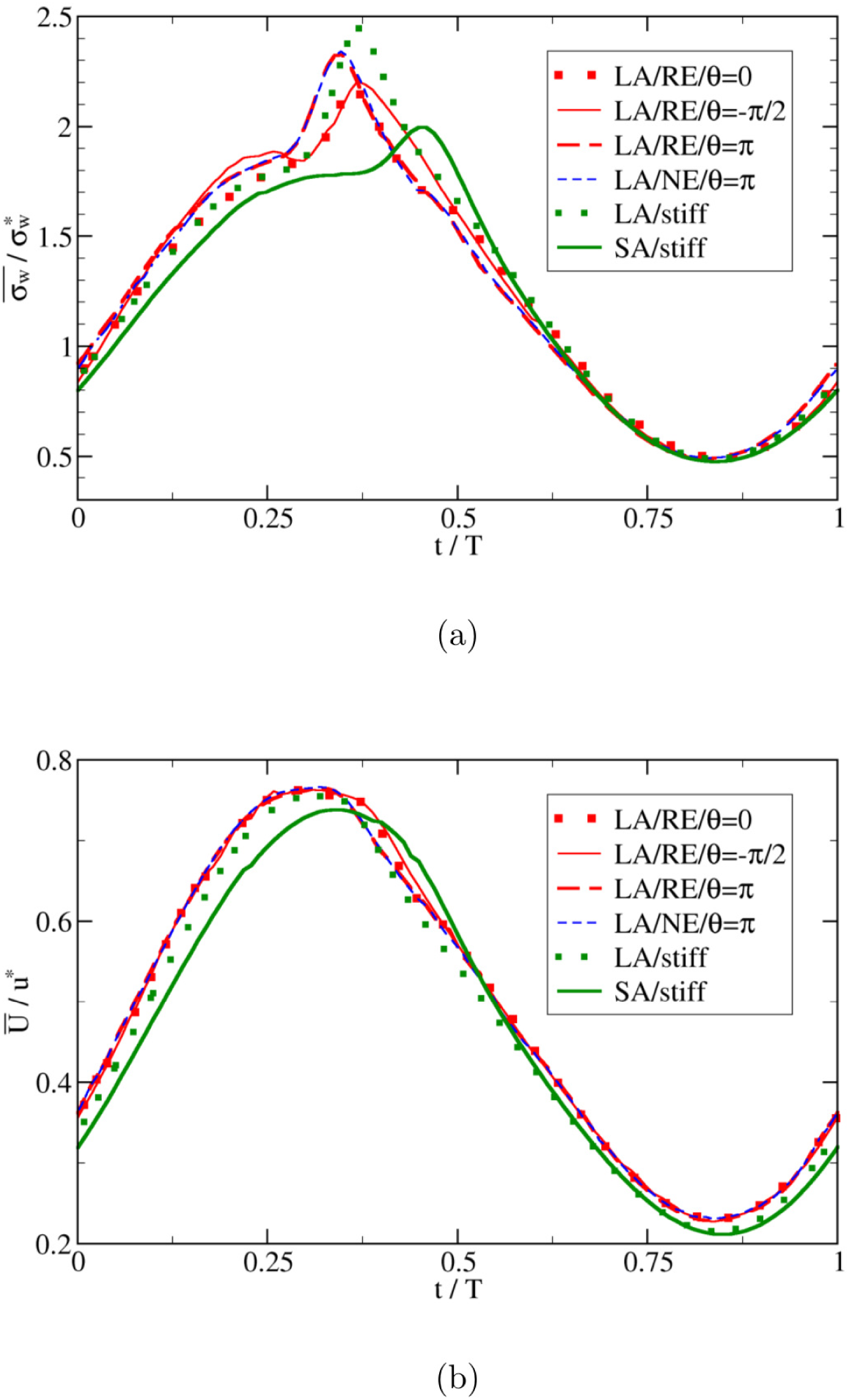
Time evolution of wall shear stress and flow velocity in parent vessel. The cross-sectional averages of (a) WSS, 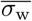, and (b) flow velocity, 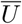, are computed in the parent vessel region (i.e., sufficiently far from the aneurysm) and normalized by their reference values in a circular tube (page 11). The flow is driven by a simple sine-type pressure gradient, *p*′. Transmural pressure, *P*_tr_, is also a sine wave but has a phase shift, *θ*, with respect to *p*′ as indicated. Acronyms in the legends: SA=Small Aneurysm; LA=Large Aneurysm; NE=Normal Elasticity (using *E*_0_) (page 12).

**Fig 11.**
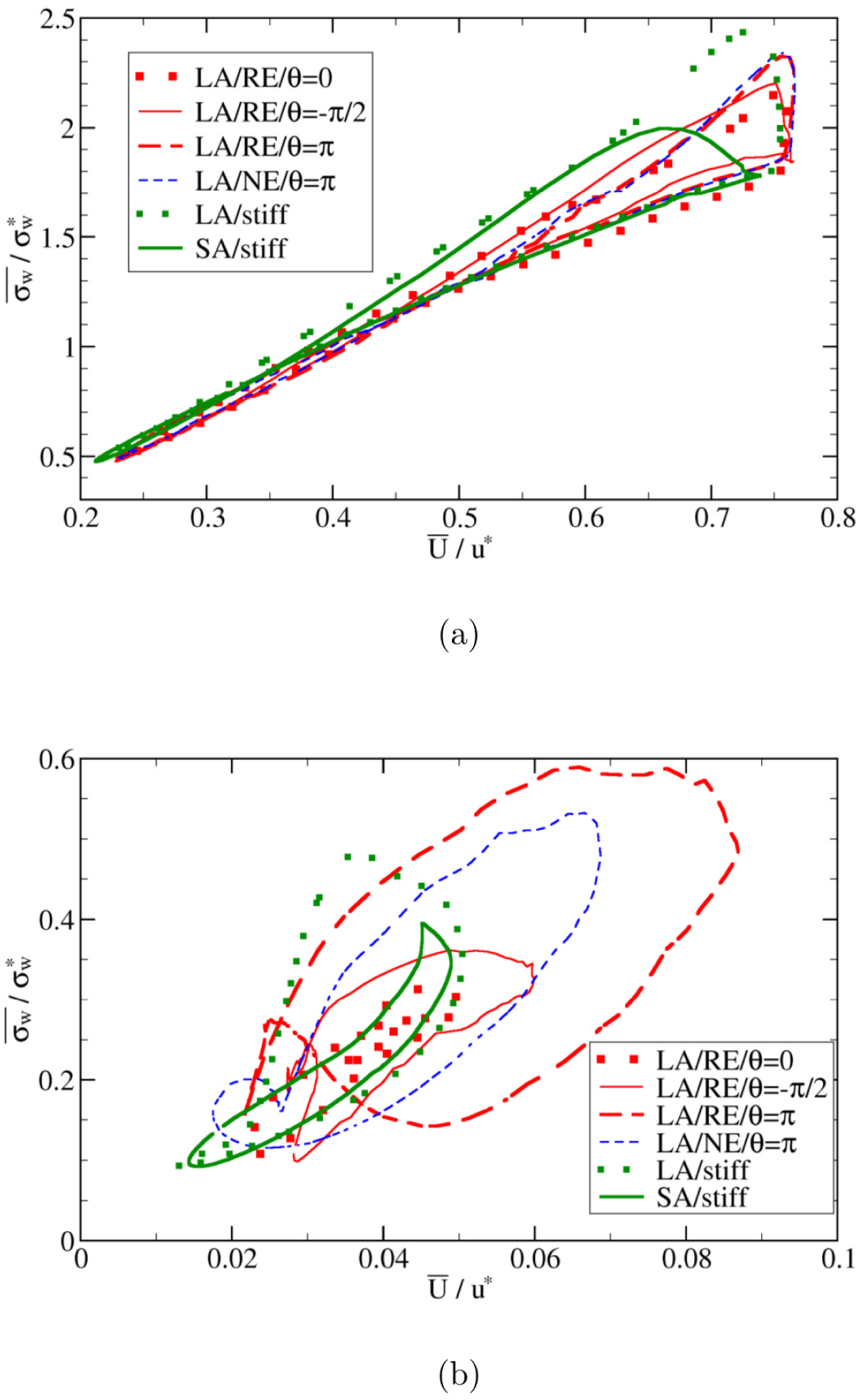
Relation between wall shear stress and flow velocity. The spatial averages of WSS, 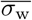, and flow velocity, 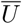, are computed (a) outside and (b) within the aneurysmal region. Acronyms in the legends: SA=Small Aneurysm; LA=Large Aneurysm; NE=Normal Elasticity (using *E*_0_); RE=Reduced Elasticity (using *E*_0_/3) (page 12). For fluid in the parent vessel, the Pearson correlation coefficients between the average flow velocity and WSS in different cases are close to unity (≥ 0.95; see text for more details). For the fluid inside the aneurysm, however, the Pearson correlation coefficients can become as small as 0.7. The WSS 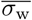 and velocity 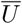 are normalized by the characteristic value 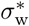 and *u*\*, respectively.

Within the aneurysmal domain, however, the flow has a more complex pattern so that a one-to-one correspondence between characteristic flow velocity and wall shear stress is no longer ensured. This is particularly the case for different values of the phase shift *θ* between pressure gradient and transmural pressure waveforms (Fig 12). From Fig 11b it is evident that the correlation between the velocity and WSS within the aneurysm is deteriorating with increasing phase shift angle *θ*. In terms of Pearson correlation coefficient, one obtains 0.91 (*θ* = 0), 0.76 (*θ* = −*π/*2), 0.67 (*θ* = *π*), 0.84 (LA/NE/*θ* = *π*), 0.72 (LA/stiff) and 0.93 (SA/stiff). One contributing factor to this deterioration is that the transmural pressure directly controls the aneurysm wall motion, which in return affects the IA flow and WSS.

**Fig 12.**
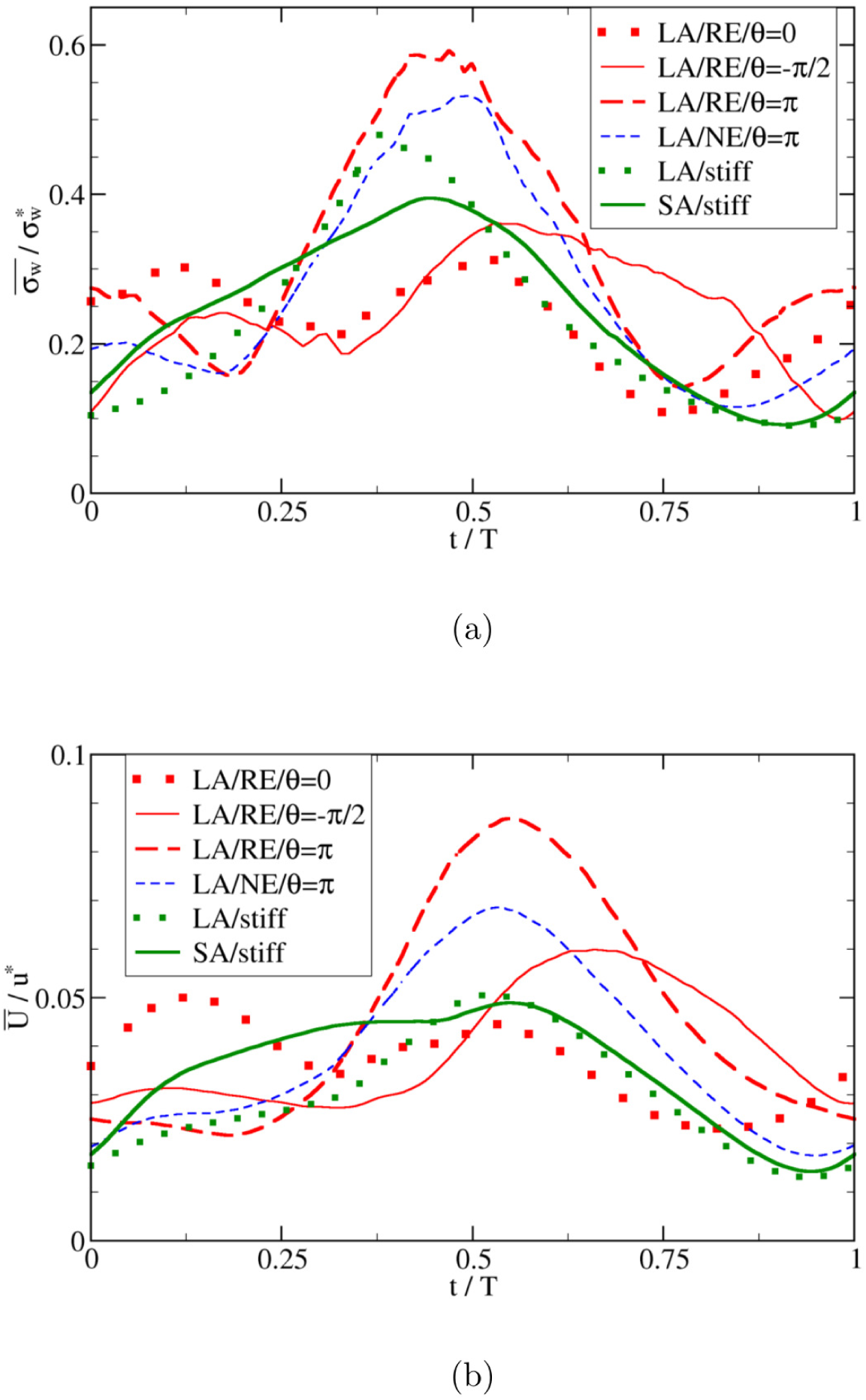
Same quantities as in Fig 10 but evaluated as spatial averages within the aneurysm dome.

Results shown in Fig 11b also reveal effects of the aneurysm size on the correlations between flow velocity and WSS. It can be seen from this data that the proxy role of flow velocity as an estimate of WSS is less justified in the case of larger and more flexible domes. This finding is of great practical importance as serious pathological cases are often associated with large aneurysm sizes and an advanced state of degradation, which leads to a weak elasticity and hence large deformability.

## Conclusion

In this work, an immersed-boundary-lattice-Boltzmann-finite-element method is employed to simulate three-dimensional viscous flow in an elastic vessel containing a side-wall aneurysm, with a special focus on the relevance of the aneurysm deformability.

Accuracy of the method is shown through steady Poiseuille and pulsatile Womersley flow tests in a circular rigid tube. Furthermore, the deformation behavior of an elastic pipe under the effect of transmural pressure is studied using the present model and validated by comparisons to both SimVascular and the tube law.

Hemodynamic quantities, such as flow velocity and wall shear stress (WSS), play a crucial role in the development and failure of aneurysms. Using the above described simulation tool, we investigate the combined effect of aneurysm size and wall softness on intra-aneurysm (IA) flow parameters. Under the rigid wall assumption, the IA flow velocity and WSS decrease with increasing aneurysm size. However, the situation becomes more complex when the aneurysm deformation becomes important.

The central result of our investigation concerns the connection between WSS and flow velocity, the latter often used as a proxy observable to estimate the former. While an indirect evaluation of the WSS via the flow velocity is a reliable method for regular geometries (planar or cylindrical), the present study reveals that the correlation between the two quantities deteriorates within the aneurysmal domain. The situation becomes more problematic in the case of large and easily deformable aneurysms.

In aneurysms, arterial wall stiffness can significantly decrease due to tissue degradation. This is often accompanied by an increase of the aneurysm’s size. Our results suggest that, in such serious pathological cases, fluid velocity is no longer a reliable proxy for WSS.

## Supporting information

**S1 Appendix. Temporal variation of fluid velocity field within a planar cut of the aneurysm.**

## Acknowledgments

HW acknowledges his IMPRS-SurMat scholarship and the financial support by the STKS department at ICAMS, Ruhr-University Bochum. TK thanks the University of Edinburgh for the award of a Chancellor’s Fellowship. We thank Arunn Sathasivam for his support in the numerical solution of Womersley flow. Useful discussions with Alessio Alexiadis are gratefully acknowledged. ICAMS acknowledges funding from its industrial sponsors, the state of North-Rhine Westphalia and the European Commission in the framework of the European Regional Development Fund (ERDF).

